# Dynamic ALFF Reveals Phase- and Network-Dependent Responses During Acute Severe Hypoxia

**DOI:** 10.64898/2026.03.27.713294

**Authors:** Daehun Kang, Koji Uchida, Clifton R. Haider, Norbert G. Campeau, Myung-Ho In, Erin M. Gray, Joshua D. Trzasko, Kirk M. Welker, Chad C. Wiggins, Jonathon W. Senefeld, Sarah E. Baker, Matt A. Bernstein, David R. Holmes, Michael J. Joyner, Timothy B. Curry, John Huston, Yunhong Shu

## Abstract

1.

Hypoxia constrains cerebral oxygen availability and challenges brain function. Previous work showed that functional connectivity reorganizes early during acute hypoxia, preceding cognitive deterioration, but the functional changes accompanying more severe hypoxic stress remain incompletely understood. We examined dynamic amplitude of low-frequency fluctuations (dALFF) in blood-oxygenation-level dependent (BOLD) fMRI during normoxia, sustained mild hypoxia, and transient severe hypoxia in healthy adults performing a continuous Go/No-go task with concurrent physiological monitoring. We characterized dALFF at whole-brain and network levels using a causal sliding-window approach, principal component analysis, and Schaefer’s 17-network parcellation. Severe hypoxia elicited a non-monotonic, phase-dependent dALFF response that was not observed during normoxia or sustained mild hypoxia. Relative preservation during early hypoxia was followed by late-hypoxia suppression, which we operationally defined as a decompensation phase, and by a pronounced rebound after reoxygenation. Within this global response, dALFF became increasingly differentiated across intrinsic brain networks: DefaultA showed marked suppression, whereas SomMotB exhibited relative preservation or enhancement during decompensation. These changes were neither spatially uniform nor tightly synchronized with systemic oxygenation, while broadly overlapping temporally with cognitive deterioration. Together, these findings indicate that dALFF captures a complementary aspect of the brain’s response to acute hypoxic stress, characterized by reversible, phase- and network-dependent reorganization of ongoing low-frequency BOLD dynamics under constrained oxygen availability.

**Highlights:**

- Severe hypoxia evokes a non-monotonic, phase-dependent dynamic ALFF response.
- dALFF changes are globally coherent yet network-specific during severe hypoxia.
- DefaultA is suppressed while SomMotB is relatively preserved during decompensation.
- This network differentiation broadly overlaps with cognitive deterioration.
- dALFF may help identify the onset of reversible decompensation during hypoxic stress.

## 2. Introduction

Oxygen is essential for maintaining cerebral metabolism (Raichle and Gusnard, 2002), and even brief reductions in its availability can trigger a wide array of physiological and cognitive alterations (McMorris et al., 2017; Post et al., 2023; Bloomfield et al., 2026). Physiologically, a decline in arterial oxygen partial pressure (P_a_O_2_) is the earliest and most sensitive indicator of hypoxic exposure (Bengtsson et al., 2001; Ortiz-Prado et al., 2019), although cerebral oxygen delivery ultimately depends on arterial oxygen content (CaO_2_) and blood flow (Todd et al., 1994; Turner et al., 2024). In response to reduced oxygen availability, the brain attempts to maintain metabolic homeostasis by increasing cerebral blood flow through vasodilation and by adjusting the rate of oxygen metabolism to sustain or prioritize energy production (Brugniaux et al., 2007; Chen et al., 2007; Harris et al., 2013; Ogoh, 2015; Kang et al., 2025; Mascarenhas et al., 2025). Although compensatory, these responses may not fully mitigate the effects of hypoxia, which can lead to temporary or permanent cognitive impairment and neural dysfunction (Phillips et al., 2015; Turner et al., 2015; McMorris et al., 2017; Naismith et al., 2020; Uchida et al., 2020a; Uchida et al., 2020b; Wang et al., 2022; Gong et al., 2025; Grzybowska-Ganszczyk et al., 2025). These effects vary with the severity and duration of hypoxic exposure and may manifest differently across physiological, behavioral, and neuroimaging measures (McMorris et al., 2017; Bloomfield et al., 2026).

Our previous work has shown that acute hypoxia-induced changes in functional connectivity (FC) correlate with reductions in end-tidal oxygen partial pressure (P_ET_O_2_), a surrogate for arterial oxygen tension (P_a_O_2_), and emerge at a physiological threshold (Kang et al., 2026). These findings revealed a proactive and structured reconfiguration of large-scale brain networks in response to hypoxic stress, characterized by a selective increase in inter-network connectivity centered on the default mode network (DMN) during hypoxia. Notably, although FC increased following threshold crossing of P_ET_O_2_, these changes did not track the timing or magnitude of behavioral deterioration. FC reorganization was observed in both severe and mild hypoxia (fraction of inspired oxygen F_i_O_2_= 7.7% and 11.8%, respectively) (Kang et al., 2026). In severe hypoxia, these changes preceded behavioral decline (Kang et al., 2026), whereas the mild hypoxia did not elicit significant behavioral deterioration in prior work using the same paradigm (Uchida et al., 2020b). This dissociation indicates that FC reconfiguration may reflect an early response to reduced oxygen availability, while the functional changes accompanying subsequent cognitive deterioration remain incompletely understood.

Amplitude of low-frequency fluctuations (ALFF), as voxel-wise metrics derived from resting-state blood-oxygenation-level dependent (BOLD) signals, has been widely used to characterize regional alterations in spontaneous brain activity across neurological and psychiatric disorders (Yang et al., 2007; Han et al., 2011; Achuthan et al., 2023; Wang et al., 2025). ALFF quantifies the power of BOLD signal fluctuations at low frequencies (0.01-0.10 Hz) (Achuthan et al., 2023; Biswal and Uddin, 2025), whereas FC characterizes the temporal coupling of the BOLD fluctuations between spatially distributed brain regions. Together, these measures capture complementary aspects of BOLD dynamics, reflecting regional signal amplitude and large-scale network coordination, respectively (Di and Biswal, 2015; Tommasin et al., 2017). A time-resolved ALFF approach may be particularly informative during acute hypoxia, where rapidly changing physiological conditions may produce transient, phase-dependent alterations in spontaneous BOLD activity that are not captured by static measures (Fu et al., 2018; Zheng et al., 2021). In addition, we examined dynamic amplitudes of high-frequency fluctuations (dAHFF, > 0.10 Hz) as a complementary measure of higher-frequency BOLD modulation that may be more sensitive to cardiac, respiratory, and vascular influences (Birn, 2012; Delli Pizzi et al., 2023).

In the present study, we hypothesized that dynamic ALFF would capture complementary aspects of the brain’s response to acute hypoxia relative to functional connectivity, and that these responses would exhibit structured variation across intrinsic brain networks. We characterized whole-brain dynamic ALFF (dALFF) and dAHFF trajectories under normoxia, sustained mild hypoxia, and transient severe hypoxia. Because significant cognitive deterioration was observed only during severe hypoxia, we focused additional analyses on this condition and its recovery period. We examined dALFF in relation to systemic oxygenation measures and cognitive deterioration indexed by commission errors in a Go/No-Go task, and applied principal component and network-level analyses to characterize its dominant temporal patterns and organization across intrinsic brain networks.

## 3. Materials and methods

### 3.1. Participants and acute hypoxia paradigm

Eleven healthy adults (five females / six males; age 26.5 ± 4.5 years, mean ± SD) were recruited for this study. All participants were self-reported as healthy, with no known vascular, respiratory, cardiac, or neurological conditions. Prior to the experimental session, participants were familiarized with the respiratory circuit and the cognitive task outside the MRI suite. Baseline physiological characteristics of the cohort reported in our previous study (Kang et al., 2026) and are not reproduced here because they were not used in the present analyses. The study was conducted under an Institutional Review Board–approved protocol (Mayo Clinic IRB#16-004354), and all participants provided written informed consent prior to participation. Experimental procedures were performed in accordance with the ethical standards set by the *Declaration of Helsinki*.

Each participant underwent three 10-min fMRI scans in the following order: normoxia, severe hypoxia, and mild hypoxia. The severe hypoxia scan followed a block design consisting of three minutes of normoxia (F_i_O_2_ = 21%) (baseline), followed by three minutes of hypoxic gas (F_i_O_2_ = 7.7%) administration, and four minutes of recovery with normoxic air. In contrast, the normoxia and mild hypoxia scans were performed under continuous delivery of the assigned gas mixture throughout the 10-min acquisition (F_i_O_2_ = 21% and 11.8%, respectively). A minimum interval of 20 min was maintained between scans. Participants were instrumented with a two-way non-rebreathing valve (2700 Series, Hans Rudolph, Shawnee, KS, USA) attached to a facemask (7450 Series V2, Hans Rudolph, Shawnee, KS, USA). During the normoxic inspirate condition, participants inspired compressed room air (∼21% oxygen balanced with nitrogen). Normobaric hypoxia inspirates consisted of pre-mixed tanks containing 11.8% O_2_ (moderate hypoxia) and 7.7% O_2_ (severe hypoxia) balanced with nitrogen. All gases were delivered from a Douglas bag reservoir containing the pre-mixed gas. During all fMRI scans, participants performed a computer-based Go/No-Go cognitive task (Uchida et al., 2020b).

To ensure subject safety, a board-certified anesthesiologist was present throughout all MRI sessions, including hypoxic challenges, continuously monitoring participant condition and cardiorespiratory parameters.

### 3.2. Data acquisition

All participants were scanned on a compact 3T MRI system (C3T) equipped with a high-performance gradient (80 mT/m peak amplitude, 700 T/m/s peak slew rate) (Foo et al., 2018), using a 32-channel head coil (Nova Medical, Wilmington, MA, USA).

Functional images were acquired using a gradient-echo echo-planar imaging (GRE-EPI) sequence during each 10-min scan (TR = 2 s, TE = 30 ms, flip angle = 90°, isotropic resolution = 2.5 mm, simultaneous multi-slice acceleration factor = 3, no in-plane acceleration, 300 volumes). A high-resolution T1-weighted anatomical image was acquired for spatial reference using a magnetization-prepared rapid acquisition with gradient echo (MPRAGE) sequence (TR = 5.4 ms, TE = 2.4 ms, TI = 1000 ms, flip angle = 8°, isotropic resolution = 1.0 mm). All MRI data were retrospectively corrected for gradient nonlinearity distortions using an inline tenth-order gradwarp algorithm (Tao et al., 2017).

Physiological signals were continuously recorded during functional imaging. Arterial blood pressure was monitored noninvasively using finger photoplethysmography (Nexfin, TPD Biomedical Instrumentation). Heart rate, peripheral oxygen saturation (S_p_O_2_), and breath-by-breath end-tidal O_2_ and CO_2_ were continuously monitored (Cardiocap/5, Datex-Ohmeda). Participants breathed through a low-dead-space, two-way non-rebreathing valve circuit, with expiratory flow measured using a turbine system (Universal Ventilation Meter, Ventura, CA). Analog signals were collected and converted to digital signals at a rate of ∼1,000 Hz using a data acquisition system (PowerLab 16/30 and LabChart 8, ADInstruments, Colorado Springs, CO, USA).

### 3.3. Cognitive and behavioral measures

During the functional imaging sessions, the participants performed a computer-based Go/No-Go cognitive task, implemented using MATLAB software (R2019b, MathWorks, Natick, MA) (Uchida et al., 2020b). Task stimuli were presented visually using an LCD display system (Nordic NeuroLab, Bergen, Norway). The subjects were instructed to press a button in response to a “Go” stimulus but refrain from pressing the button for a “No-Go” stimulus.

Behavioral performance was quantified using commission error rate, omission error rate, and reaction time. Commission errors were defined as incorrect responses to No-Go stimuli, whereas omission errors reflected failures to respond to Go stimuli. Reaction time was defined as the interval between stimulus presentation and button press. These metrics were continuously recorded throughout the task. To ensure adequate task familiarization, participants completed a 10-min practice session prior to scanning, as previously recommended (Taylor et al., 2015). Based on prior evidence that commission error rates increase under acute hypoxia and sensitively reflect impaired inhibitory control, commission error count was selected as the primary behavioral marker of hypoxia-induced cognitive deterioration (Uchida et al., 2020b).

### 3.4. Data preprocessing and signal derivation

#### 3.4.1. Preprocessing for MR images

All functional MRI data, consisting of 300 consecutive EPI volumes per scan, were preprocessed. Preprocessing steps included removal of the first 10 volumes, de-spiking, slice-timing correction, physiological noise correction using RETROICOR (Glover et al., 2000; Birn et al., 2008), EPI–MPRAGE alignment, inter-volume motion correction, and spatial smoothing with a Gaussian kernel (4 mm full-width at half-maximum, FWHM). Signal intensities were then scaled to a mean value of 100.

Subsequently, nuisance regression was performed using linear regressors including Legendre polynomials (up to 4th order), rigid-body motion parameters, and white matter and cerebrospinal fluid (WM/CSF) signals derived using ANATICOR and CompCor approaches (Behzadi et al., 2007; Jo et al., 2010). Residual BOLD time series were obtained following regression using the AFNI software package (Cox, 1996). Motion censoring was not applied because removal of individual volumes would disrupt the continuous time series required for sliding-window spectral analysis.

T1-weighted anatomical images were processed to segment brain structures using FreeSurfer (Reuter et al., 2012). The resulting anatomical segmentations were used for coregistration to individual EPI images and for projection onto standard cortical surfaces provided by AFNI and SUMA (Cox, 1996).

#### 3.4.2. Pre-defined ROI with intrinsic brain networks

Regions of interest (ROIs) were defined using the 400-region cortical parcellation from the Schaefer atlas (Schaefer et al., 2018). The atlas was implemented on the AFNI standard surface model (https://afni.nimh.nih.gov/pub/dist/atlases/SchaeferYeo/) and subsequently transformed into individual subjects’ native volumetric space for analysis of EPI data. Each ROI was assigned to one of 17 intrinsic connectivity networks, enabling analyses at both the ROI and network levels for functional connectivity and spontaneous BOLD signal fluctuations.

#### 3.4.3. Dynamic changes by sliding window

To examine dynamic ALFF (dALFF), mean BOLD time series were extracted from each of the 400 ROIs using the preprocessed residual BOLD signals. ALFF was computed for each ROI using a fast Fourier transform (FFT function) by calculating the square root of the power spectrum within the low-frequency band (0.01 ≤ *f* ≤ 0.10 Hz). dALFF was estimated using a causal sliding window of 90 seconds with a step size of 2 seconds, resulting in 246 overlapping windows per scan. This procedure generated a dALFF time series for each ROI in each subject. In addition, dAHFF was computed within the frequency band of 0.10 < *f* ≤ 0.25 Hz using the same FFT-based and sliding-window procedures, with the upper bound constrained by the Nyquist frequency determined by the repetition time (TR = 2 s). dALFF and dAHFF were quantified as relative changes (i.e., percent changes) with respect to a region-specific temporal baseline. A 90-s sliding window was used in the main analysis as a compromise between temporal sensitivity and stability.

Similar ALFF response patterns were observed when using alternative window lengths (120 s), indicating that the results were robust to window size (Supplementary Fig. 4a). As a sensitivity analysis, the further analyses were repeated using a 120-s causal sliding window with otherwise identical procedures (Supplementary Fig. 4).

### 3.5. Network-level and statistical analyses

#### 3.5.1. Temporal alignment and normalization of multimodal signals

To facilitate direct comparison of temporal trajectories across physiological, neuroimaging, and behavioral measures under acute severe hypoxia, aggregate time series of P_ET_O_2_, dALFF, dAHFF and behavioral error rates were z-score normalized and lightly temporally smoothed using a 5-point moving average filter (10 s window given the 2-s sampling interval), implemented with the movmean function in MATLAB. For each metric, baseline values were defined using the pre-hypoxia period (160 < t < 180 s), and all time courses were aligned such that the baseline mean was set to zero. Temporal smoothing was applied to reduce high-frequency noise while preserving low-frequency trends relevant to hypoxia-induced dynamics. This procedure facilitated visualization and comparison of the relative timing and progression of physiological, imaging-derived, and behavioral responses to hypoxia.

#### 3.5.2. Principal component analysis

To identify dALFF patterns associated with hypoxia, we applied principal component analysis (PCA) to the fMRI session that followed a block-design paradigm (normoxia–severe hypoxia–normoxia). This was performed because its structured temporal profile facilitates extraction of hypoxia-related signal patterns.

Prior to PCA, dALFF time series were z-scored within each ROI for each subject to remove subject-specific mean and variance differences. PCA was performed in MATLAB (R2019b) using default settings. The input matrix consisted of 2,706 observations × 400 ROIs, constructed by concatenating dALFF time points from 246 causal sliding windows across 11 subjects (246 × 11 = 2,706). In this formulation, each time point (window) represented one observation and each ROI was treated as a variable. Thus, PCA identified spatial patterns of ROIs whose dALFF values covaried across time and subjects. Components were ranked by the proportion of explained variance, and the dominant components were carried forward for subsequent analyses. For visualization of temporal dynamics, component time courses were then reshaped into a 246 (windows) × 11 (subjects) matrix to enable visualization and comparison of time courses across individuals and phases.

#### 3.5.3. Permutation test of network-specific organizations of dALFF PCA coefficients

To assess whether network-level dALFF PCA coefficients deviated from the overall ROI distribution beyond random variation, a permutation-based test was performed. In each permutation, dALFF PCA coefficients were randomly shuffled across ROIs while preserving the original number of ROIs within each intrinsic network. Network-wise median coefficients were recomputed using the same procedures applied to the observed data.

This process was repeated 10,000 times to generate a null distribution of median coefficients for each network. The test statistic was defined as the absolute difference between the network-wise median coefficient and the median coefficient across all ROIs. Two-sided empirical P values were calculated as *(k+1)/(N+1)*, where *k* was the number of permutations producing a deviation from the overall ROI median greater than or equal to the observed deviation, and *N* was the total number of permutations. P values across networks were corrected for multiple comparisons using the false discovery rate procedure (MATLAB’s *mafdr*). This approach tests whether the spatial distribution of dALFF PCA coefficients was organized by intrinsic network beyond that expected from random ROI assignment.

## 4. Results

### 4.1. Temporal profiles of whole-brain dALFF and dAHFF across normoxic and hypoxic conditions

Figure 1 summarizes the experimental protocols and whole-brain dALFF and dAHFF trajectories under normoxia, sustained mild hypoxia, and transient severe hypoxia. Under normoxia, dALFF gradually increased, whereas AHFF remained relatively stable (Figs. 1a, d and g). During sustained mild hypoxia (FiO2 = 11.8%), which was maintained throughout the 10-min fMRI acquisition, both measures showed only gradual temporal variation without a pronounced transient shift (Figs. 1b, e and h).

**Figure 1.**
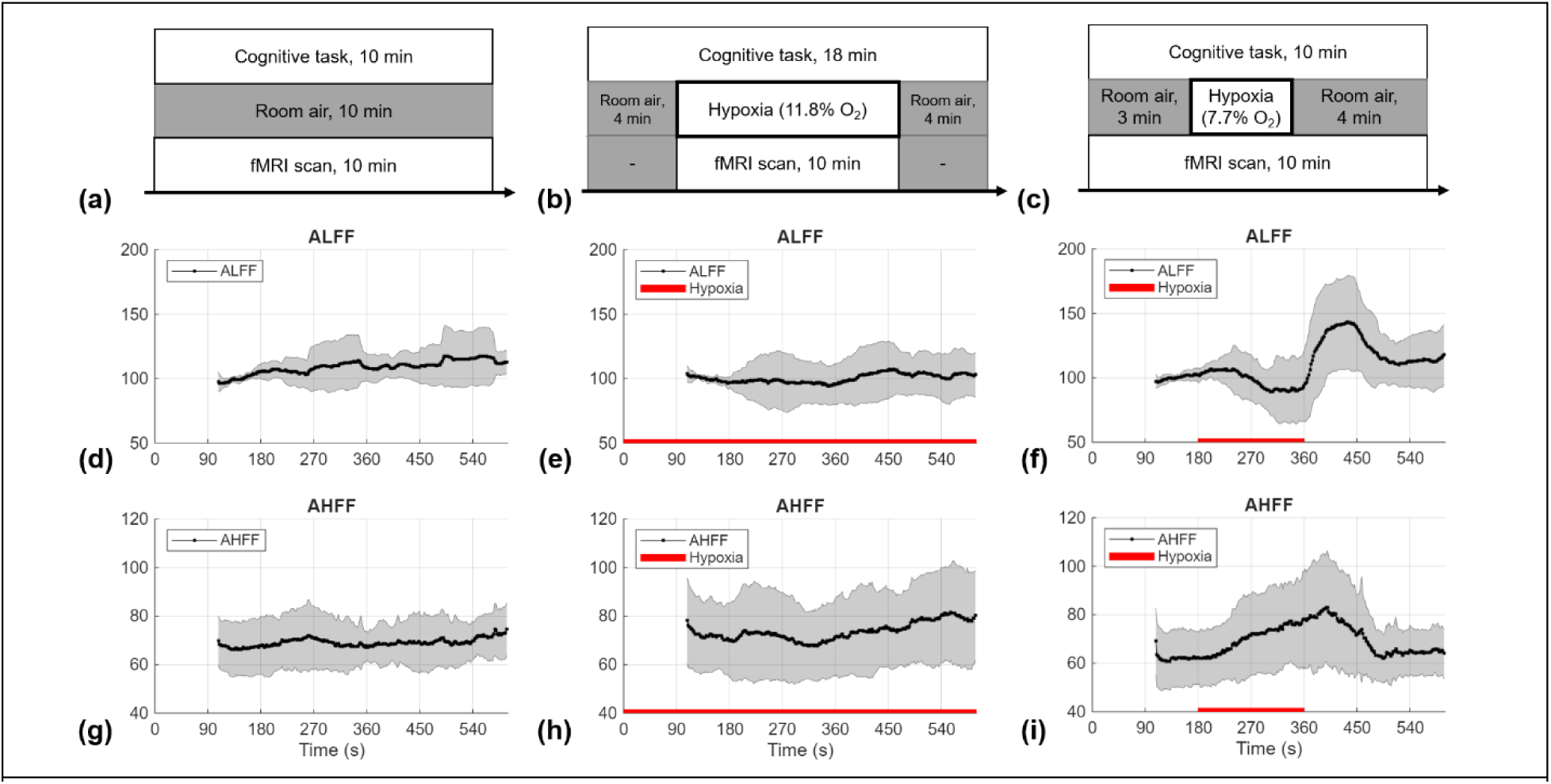
Experimental protocols and condition-specific temporal dynamics of ALFF and AHFF during normoxia, sustained mild hypoxia, and severe hypoxia. (a-c) Experimental protocols for the normoxic, mild-hypoxia, and severe-hypoxia conditions, respectively. (d-f) Group-level temporal trajectories of the amplitude of low-frequency fluctuations (ALFF; 0.01–0.10 Hz), calculated using a 90-s sliding window. (g-i) Group-level temporal trajectories of the amplitude of high-frequency fluctuations (AHFF; 0.10–0.25 Hz), calculated using a 90-s sliding window. For each participant, both ALFF and AHFF were expressed as percentages of the mean ALFF over the reference interval of 160 s < t < 180 s. Solid black lines represent the group means, and shaded regions represent ±1 standard deviation across participants in panels (d-i). Red horizontal lines in panels (e), (f), (h), and (i) indicate the hypoxic exposure periods.

Severe hypoxia (FiO2 = 7.7%; t = 180-360 s) produced more prominent changes (Fig. 1c). dALFF declined during late hypoxia and showed a marked rebound after the return to room air (Fig. 1f). In contrast, dAHFF increased during the hypoxia and declined during recovery (Fig. 1i). The temporal and network-specific dALFF responses to severe hypoxia were examined in subsequent analyses.

The late-hypoxia reduction in whole-brain dALFF during severe hypoxia also showed substantial inter-individual variability (Supplementary Fig. 1). When dALFF change between t = 180 s and t = 360 s was compared between severe hypoxia and normoxia, severe hypoxia showed a greater reduction at the group level, although the difference did not reach statistical significance (paired Wilcoxon signed-rank test, p = 0.067).

### 4.2. Multimodal temporal responses during acute severe hypoxia and recovery

We characterized the temporally aligned physiological, behavioral, and fMRI responses during a 3-min acute severe hypoxia challenge (Fig. 2). End-tidal O_2_ (P_ET_O_2_) exhibited the earliest and most pronounced change at hypoxia onset and during recovery (Fig. 2b), decreasing from a baseline of 109.9 ± 4.7 mmHg to 35.3 ± 1.7 mmHg immediately prior to reoxygenation. In contrast, SpO_2_ showed a slower temporal response (Fig. 2c), declining from 98.3 ± 1.3% to 69.3 ± 8.7%. Behavioral performance, indexed by the number of commission errors, exhibited an even more delayed response following hypoxia onset (Fig. 2d).

**Figure 2.**
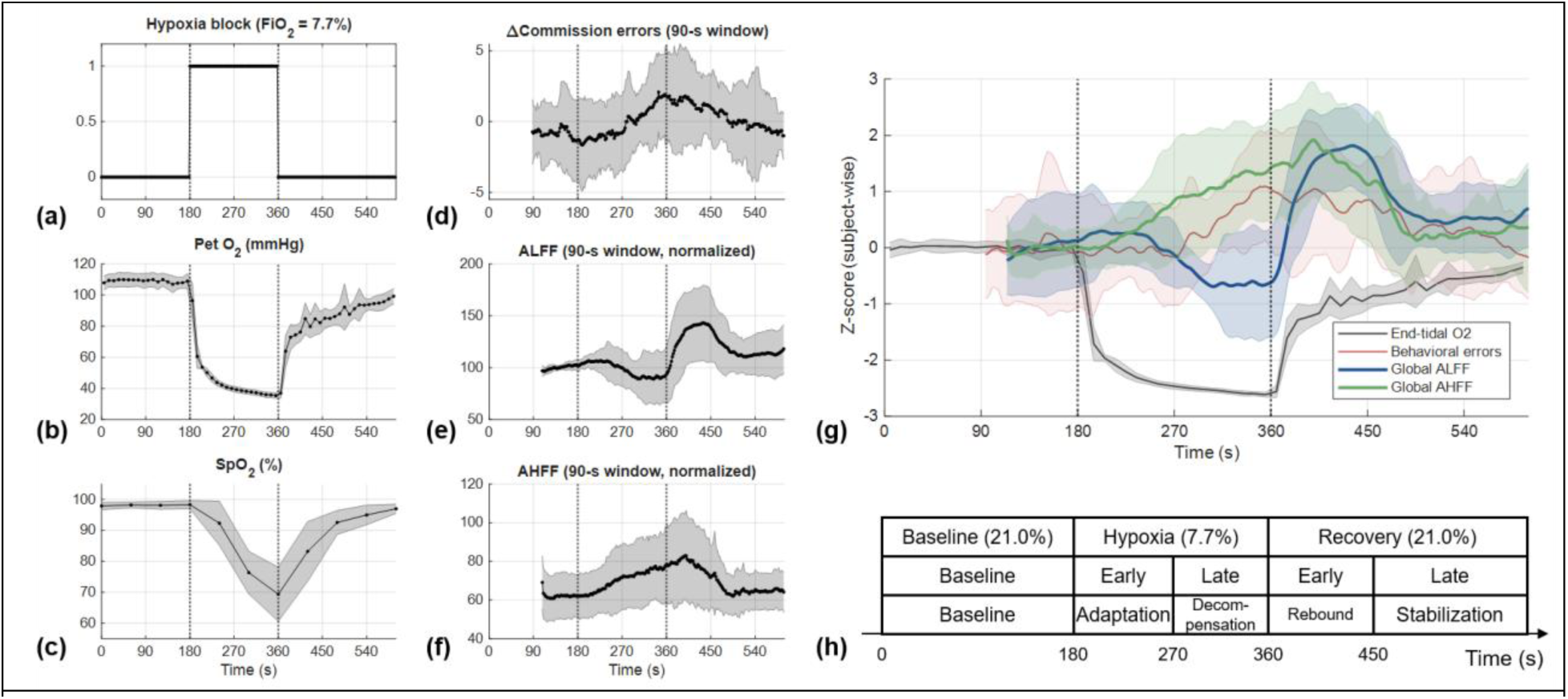
Phase-structured multimodal responses to an acute severe hypoxia challenge (F_i_O_2_ = 7.7%). (a) Experimental timing of hypoxic stimulation. (b) End-tidal oxygen partial pressure (P_ET_O_2_). (c) Peripheral oxygen saturation (SpO_2_). (d) Commission errors as behavioral deterioration. (e) Amplitude of low-frequency fluctuations (ALFF; 0.01-0.1Hz), normalized to Baseline values (t < 180 s). (f) Amplitude of high-frequency fluctuations (AHFF; 0.1-0.25Hz), normalized to Baseline values of ALFF. Panels (d-f) were estimated using a causal sliding-window approach (90 s window length). (g) Z-score normalized and lightly temporally smoothed summary of P_ET_O_2_, ALFF, AHFF and commission errors, illustrating their distinct temporal evolution. Averages of baseline values (t < 180 s) were aligned to zero. Vertical dashed lines indicate severe hypoxia onset and offset. (h) Conceptual segmentation of the experiment into baseline, hypoxia, and recovery periods, with hypoxia and recovery further divided into adaptation, decompensation, rebound, and stabilization phases based on dominant multimodal features observed across measures. Shaded regions represent ± standard deviation in panels (a-g) across subjects.

Dynamic ALFF and AHFF derived from the concurrently acquired fMRI BOLD time courses exhibited distinct temporal profiles (Fig. 2e and f). dAHFF showed a delayed increase during hypoxia, followed by a delayed decline during recovery. In contrast, dALFF exhibited a non-monotonic temporal pattern, declining during late hypoxia and showing a marked rebound after reoxygenation.

Figure 2g presents the temporally aligned and z-score-normalized trajectories of P_ET_O_2_, ALFF, AHFF and behavioral error. For descriptive comparison, the time course was divided into four phases: early hypoxia, late hypoxia, early recovery, and late recovery (Fig. 2h). Operationally, these phases correspond to adaptation (no measurable increase in ALFF and behavioral error with initial AHFF emergence), decompensation (progressive reduction in ALFF accompanied by increasing behavioral error during sustained severe hypoxia), rebound (transient overshoot in ALFF following reoxygenation), and stabilization (return toward baseline dynamics of ALFF, AHFF and behavioral error).

To further examine the relationship between systemic oxygenation and dALFF, dALFF trajectories were plotted against corresponding P_ET_O_2_ and S_p_O_2_ values under severe and mild hypoxia (Supplementary Figs. 2 and 3, respectively). During severe hypoxia, dALFF showed a phase-dependent pattern, with relative preservation during early hypoxia, progressive suppression during late hypoxia, and a marked rebound after reoxygenation, in contrast to the more monotonic changes in P_ET_O2 and S_p_O_2_. During mild hypoxia, dALFF showed a modest increase as P_ET_O_2_ approached approximately 52 mmHg and S_p_O_2_ approximately 84%; however, comparable P_ET_O_2_ and S_p_O_2_ ranges during severe hypoxia were associated with a different dALFF response. These observations indicate that similar systemic oxygenation levels can be associated with distinct dALFF dynamics across hypoxic conditions.

### 4.3. Dynamic ALFF modulation is global yet exhibits network-specific structure

To characterize the spatial organization of dynamic ALFF (dALFF) responses to acute severe hypoxia, PCA was applied to the normalized dALFF time series from all 400 ROIs. As shown in Fig. 3a, the first principal component (PC1) accounted for 52.90% of the total variance, whereas PC2 accounted for 6.17%. Together, these two components explained 59.08% of the total variance, whereas each remaining component accounted for less than 3.40%, highlighting the dominant contribution of PC1 and a secondary, but meaningful contribution of PC2.

**Figure 3.**
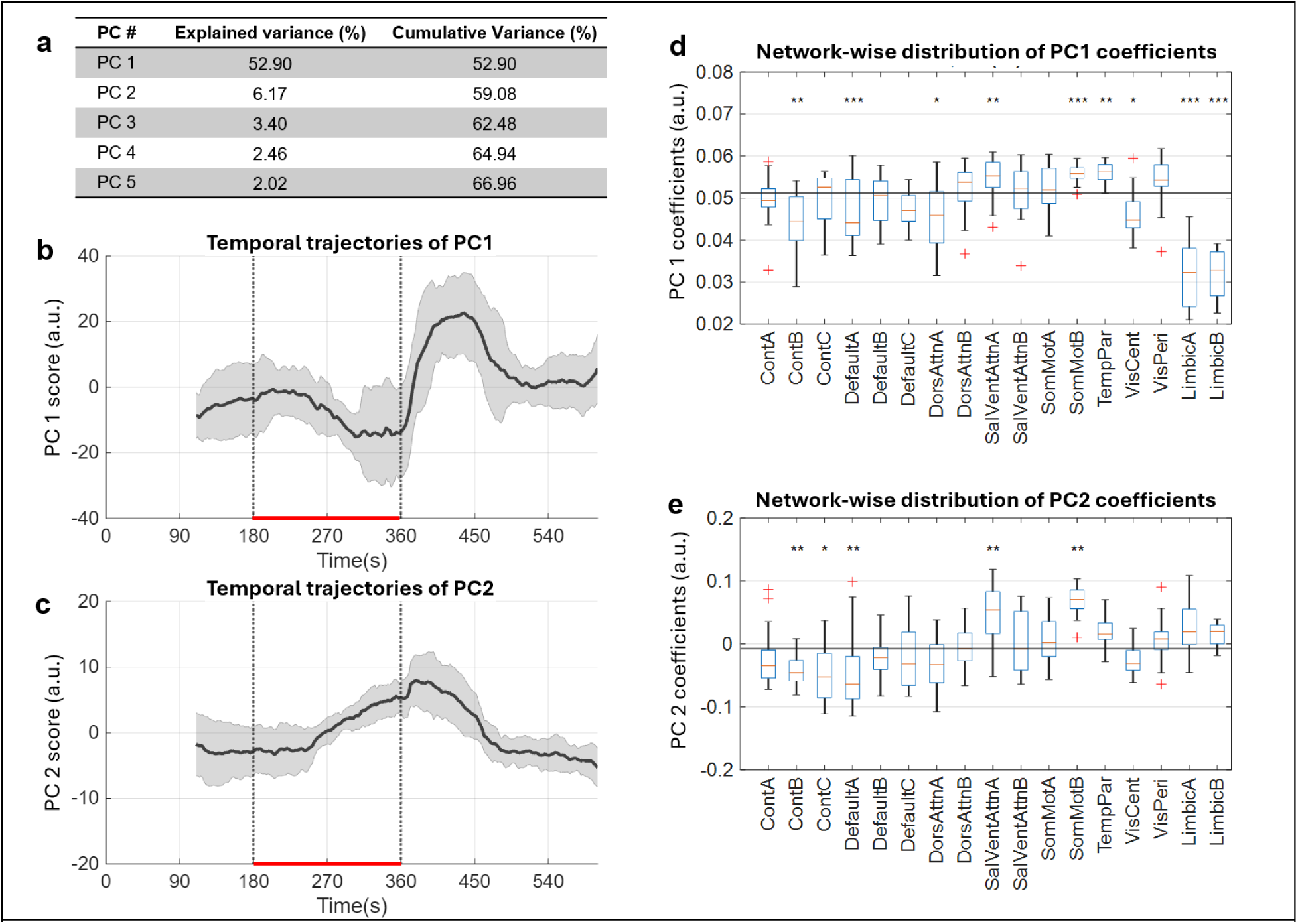
Global, non-monotonic, and network-specific dALFF responses during acute severe hypoxia. (a) Explained and cumulative variances of the first five principal components (PCs) derived from z-score–normalized ROI-level dALFF time series. (b,c) Temporal profiles of the first and second principal component (PC1 and PC2, respectively), capturing the phase-structured ALFF modulation across the experiment. (d,e) Distribution of ROI-level PC1 and PC2 coefficients across the 17 brain networks of the Schaefer atlas, illustrating network-specific contributions to the dominant ALFF mode. For each network, the observed median coefficient was compared with its permutation-derived null distribution. Asterisks indicate significant deviations from the null after correction for multiple comparisons. In panels (b, c), shaded regions represent ± standard deviation across subjects, vertical dashed lines indicate hypoxia onset and offset, and red horizontal bars denote the hypoxia period.

The temporal profile of PC1 closely resembled the global dALFF trajectory shown in Fig. 1f (Fig. 3b). All PC1 coefficients had the same sign, indicating a spatially coherent and globally synchronized ALFF modulation rather than localized or opposing effects. However, the magnitude of these coefficients varied across 17 networks, defined by the Schaefer atlas (Fig. 3d). Permutation testing showed the median PC1 coefficients were lower than expected in DefaultA, ContB, DorsAttnA, VisCent, LimbicA and LimbicB, and higher than expected in SomMotB, SalVentAttnA and TempPar. The median coefficient across all 400 ROIs was 0.0512 and is shown as a horizontal line in Fig. 3d. Interpretation of LimbicA and LimbicB was limited by signal loss in susceptibility-prone regions; these networks were therefore excluded from subsequent network comparisons.

Although PC2 explained a smaller proportion of variance (6.17%), it captured temporally specific fluctuations that were most prominent during the late hypoxia and early recovery (Fig. 3c). In contrast to PC1, PC2 included both positive and negative coefficients, indicating a more differentiated spatial pattern across networks (Fig. 3e). Median coefficients were lower than expected in DefaultA, ContB, and ContC, and higher than expected in SomMotB and SalVentAttnA relative to the overall ROI median of −0.0074.

Across both components, DefaultA had the lowest median coefficient and SomMotB the highest among the networks retained for subsequent analyses.

### 4.4. Divergent dALFF responses in DefaultA and SomMotB during severe hypoxia

To examine whether the network heterogeneity identified by PCA was evident at the subject level, dALFF changes were averaged across ROIs within DefaultA and SomMotB for each participant. Time-resolved comparisons between the two networks showed that a nominal difference (*p* < 0.05, paired Wilcoxon signed-rank test) emerged at approximately 306 s and persisted until approximately 470 s (Fig. 4a,b). This interval spanned the late-hypoxia and early-recovery periods and broadly overlapped with the period of increased Δcommission errors (Fig. 4c).

**Figure 4.**
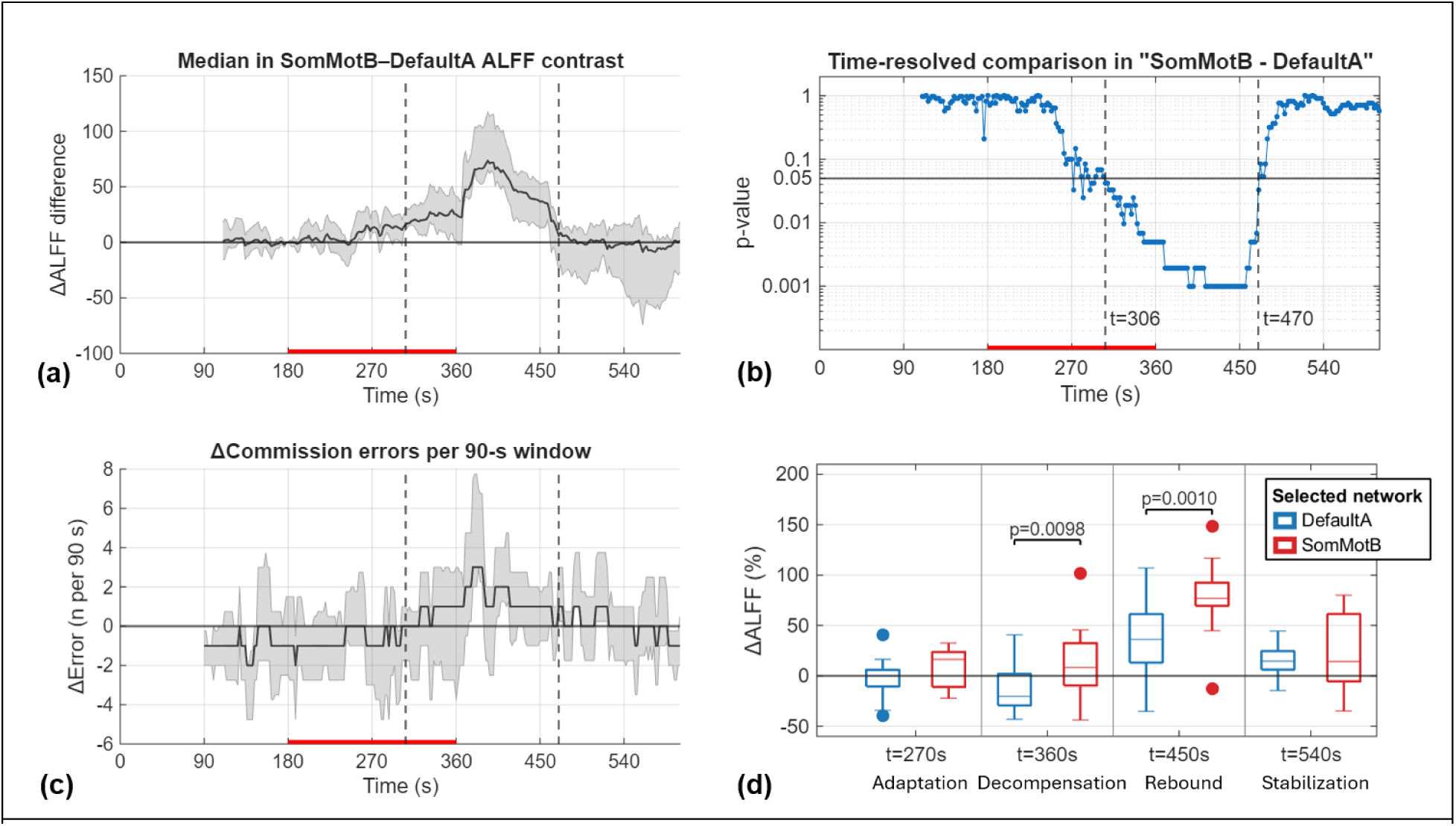
Temporal and phase-dependent differentiation in dynamic ALFF between DefaultA and SomMotB during severe hypoxia. (a) Median difference in normalized ALFF change between SomMotB and DefaultA over time, with positive values indicating relatively greater ALFF in SomMotB. (b) Time-resolved exact paired Wilcoxon signed-rank comparisons between the two networks, shown on a logarithmic p-value scale. The horizontal line indicates the nominal threshold of p = 0.05. Continuous nominal differentiation emerged at approximately 306 s and persisted until approximately 470 s. Because adjacent windows overlapped and the time-resolved tests were not corrected for multiple comparisons, this interval should be considered exploratory. (c) Median Δcommission errors across subjects, calculated using 90-s sliding windows. The period of increased Δcommission errors broadly overlapped with the interval of network differentiation. (d) Distributions of ALFF changes (ΔALFF) in DefaultA and SomMotB at representative phases of the severe-hypoxia experiment, calculated using a 90-s window length. ΔALFF is expressed as the percentage change relative to baseline (t = 160–180 s). The selected time points represent adaptation (t = 270 s), decompensation (t = 360 s), rebound (t = 450 s), and stabilization (t = 540 s). DefaultA and SomMotB showed significantly different responses during decompensation and rebound, based on exact paired Wilcoxon signed-rank tests. Boxes indicate the interquartile range, center lines indicate medians, whiskers indicate non-outlier ranges, and circles indicate outliers. Displayed p-values are unadjusted. Red bars in panels (a-c) denote the hypoxia exposure period. In panels (a) and (c), solid lines indicate medians and shaded regions indicate interquartile ranges (IQRs).

To further characterize the magnitude and phase dependence of this network divergence, dALFF values were summarized as medians [Q1, Q3] at four representative time points corresponding to adaptation, decompensation, rebound, and stabilization (Fig. 4d). During the adaptation phase (t = 270 s), dALFF was 0.1% [-10.6%, 6.0%] in DefaultA and 16.4% [-11.0%, 23.6%] in SomMotB, with no significant between-network difference.

During the decompensation phase (t = 360s), dALFF decreased to −20.3% [-29.4%, 2.0%] in DefaultA but remained at 8.4% [-9.5%, 32.4%] in SomMotB. The two networks differed significantly (paired Wilcoxon signed-rank test, p = 0.0098), indicating network-level divergence during late hypoxia.

During the rebound phase (t = 450s), dALFF increased to 36.3% [13.2%, 61.3%] in DefaultA and 76.8% [69.5%, 92.4%] in SomMotB. Despite the widespread post-hypoxia increase following reoxygenation, the between-network difference remained significant (p = 0.0010). During the stabilization phase (t = 540s), dALFF approached the reference level in both DefaultA (14.6% [6.2%, 24.5%]) and SomMotB (14.2% [-5.4%, 61.4%]), with no significant difference between networks.

To assess the robustness of the temporal differentiation to sliding-window length, the analysis was repeated using a 120-s window (Supplementary Fig. 4). A broadly similar pattern was observed, with nominal DefaultA-SomMotB differentiation extending from approximately 338 s to 488 s (Supplementary Fig. 4b,c) and again broadly overlapping with the period of increased Δcommission errors (Supplementary Fig. 4d). Although the exact temporal extent shifted modestly with window length, the overall phase-dependent pattern of network differentiation was preserved. Since adjacent sliding windows were highly overlapping and the repeated time-resolved comparisons were not corrected for multiple comparisons, the precise onset and duration of the nominally significant interval should be interpreted cautiously.

ROI-level inspection further showed that the DefaultA reduction was distributed across multiple posterior cingulate/precuneus and inferior parietal ROIs, whereas the largest positive median changes within SomMotB were observed primarily in auditory, insular, and secondary somatosensory ROIs (Supplementary Tables 1 and 2).

## 5. Discussion

In this study, we investigated hypoxia-induced dynamic changes in spontaneous BOLD activity using dALFF and interpreted these responses within the framework of canonical large-scale brain networks. Our findings revealed that under severe hypoxia accompanied by cognitive deterioration and subsequent recovery, dALFF exhibited distinct temporal profiles, whereas comparable changes were not observed during normoxia or mild hypoxia. In particular, divergent responses between DefaultA and SomMotB emerged during the late-hypoxia phase and persisted into early recovery. The divergence suggests that dALFF captures a complementary layer of hypoxia-induced functional dynamics characterized by network-specific responses. Importantly, the emergence of this network-level divergence broadly coincided with the period of cognitive deterioration.

The temporal behavior of dALFF across the three experimental conditions suggests that its response to hypoxic stress is not simply proportional to the duration or magnitude of oxygen reduction. Whereas normoxia and sustained mild hypoxia showed relatively gradual temporal variation, even a brief severe hypoxic challenge elicited a marked phase-dependent response, including late-hypoxia suppression and a prominent rebound following reoxygenation. Although the mild- and severe-hypoxia paradigms differed in temporal structure and therefore do not constitute a direct dose-response comparison, the distinct dALFF trajectories observed at comparable P_ET_O_2_ and S_p_O_2_ levels suggest a non-monotonic, state-dependent response to hypoxic stress. This pattern differs from the previously reported hypoxia-induced functional connectivity response, which emerged near a critical P_ET_O_2_ threshold (Kang et al., 2026), suggesting that dALFF does not simply track a single systemic oxygenation signal.

During severe hypoxia, the late-hypoxia period was distinguished from the preceding adaptation phase by suppression of dALFF and the emergence of cognitive deterioration, although the magnitude of the ALFF response varied substantially across individuals. We therefore operationally refer to this interval as the decompensation phase. Importantly, the marked dALFF rebound and recovery of cognitive performance following reoxygenation indicate that this decompensated state was reversible rather than persistently suppressed. Together with the lack of tight coupling between dALFF and either P_ET_O_2_ or S_p_O_2_, this reversibility raises the possibility that late-hypoxia dALFF suppression reflects a regulated, state-dependent adjustment of spontaneous BOLD dynamics rather than a purely passive consequence of reduced systemic oxygenation.

Within this global phase-dependent response, dALFF became progressively differentiated across intrinsic brain networks, with the most pronounced divergence emerging during the decompensation phase and extending into the rebound period. This interval also broadly overlapped with the period of cognitive deterioration. However, the magnitude of differentiation between SomMotB and DefaultA at the end of the decompensation phase (t = 360 s) showed essentially no association with commission errors across individuals (r = −0.05; data not shown), arguing against a simple model in which this network-level divergence directly determines behavioral impairment. Given that Go/No-go performance depends on distributed cognitive processes rather than a single network pair, the observed differentiation may instead reflect a broader transition in the functional state of the brain under severe hypoxic stress. Notably, DefaultA and SomMotB were also prominent components of the hypoxia-responsive functional connectivity pattern in our previous study (Kang et al., 2026), as evident in its network-level analyses. Their prominence across complementary BOLD measures suggests that these networks are consistently involved in the brain’s response to acute hypoxic stress, despite exhibiting markedly different local amplitude dynamics.

This interpretation is particularly relevant to DefaultA, a subdivision of the default mode network (DMN). The DMN is characterized by high intrinsic metabolic activity and plays a central role in the brain’s ongoing baseline functional state rather than being dedicated to a specific task operation (Raichle, 2015). Given the substantial energetic demands associated with spontaneous brain activity, the pronounced reduction in DefaultA ALFF during the decompensation phase may reflect downregulation or perturbation of this intrinsic functional state under severe metabolic constraint. Such suppression may be consistent with reduced engagement of energetically costly ongoing processes under hypoxic stress (Hofmeijer et al., 2014), although the present data do not directly resolve the underlying neuronal or synaptic mechanisms.

In contrast, SomMotB showed relative preservation or enhancement of ALFF during the hypoxic decompensation phase. In the 17-network parcellation (Yeo et al., 2011), SomMotB comprises ventral somatomotor regions including tongue-related areas, auditory cortex, and the secondary somatosensory cortex (S2), in contrast to the dorsal subnetwork SomMotA, which is more closely associated with externally oriented sensorimotor processing. The relative prominence of right-hemisphere auditory, insular, and S2 ROIs (supplementary table 2) is notable given the proposed role of this region in interoceptive and viscerosensory processing (Craig, 2009). In the present experimental context, hypoxia represents an internal physiological stress rather than a salient external sensory threat. One possible interpretation is therefore that networks involved in processing or monitoring internal bodily states remain relatively engaged during severe hypoxic stress. Whether the relative preservation of SomMotB ALFF reflects sustained interoceptive or viscerosensory processing requires further investigation.

The dALFF changes observed during the late-hypoxia phase paralleled cerebral metabolic rate of oxygen (CMRO₂) changes previously reported during severe hypoxia in the same subjects (Kang et al., 2025). After matching ROI-level results across the two analyses and aggregating them at the network level, similar spatial modulation patterns were observed, with network-level ALFF and CMRO_2_ changes showing a moderate correspondence (R^2^ = 0.37; Supplementary Fig. 5). Although exploratory, this relationship suggests that network-level dALFF modulation during severe hypoxia may be related to concurrent changes in cerebral oxidative metabolism.

Taken together, these findings suggest that acute severe hypoxia induces a reversible transition in spontaneous BOLD dynamics that is both phase dependent and spatially organized across intrinsic brain networks. The emergence of network differentiation during decompensation, its persistence into rebound, and its partial correspondence with previously observed metabolic changes indicate that dALFF captures a distinct aspect of the brain’s response to metabolic stress. Since these changes were neither spatially uniform nor tightly synchronized with systemic oxygenation, they are not readily explained as a purely passive consequence of reduced oxygen availability. Rather, the observed pattern is more consistent with a structured reorganization of spontaneous BOLD dynamics during severe hypoxic stress.

Several limitations should be acknowledged. First, because ALFF is derived from the BOLD signal, the observed changes may reflect a combination of neural, vascular, and systemic physiological influences, particularly under hypoxic conditions. Although P_ET_CO_2_ decreased during hypoxia in the study (data not shown), making a simple hypercapnia-related vasodilatory explanation unlikely, the relative contributions of neural and non-neural factors cannot be fully disentangled in the present study. Accordingly, dALFF should be interpreted as a measure of spontaneous BOLD dynamics rather than a direct measure of neuronal activity or metabolism. Second, the sample size was modest, and the findings therefore require replication in larger cohorts. Nevertheless, the within-subject graded hypoxia design and PCA-based identification of dominant temporal patterns enabled detection of coherent group-level responses despite substantial inter-individual variability. Third, network-level analysis based on the Schaefer parcellation enabled characterization of large-scale organizational patterns while necessarily averaging over regional variability within individual networks. ROI-level descriptive analyses confirmed that the magnitude and direction of dALFF changes varied across constituent regions of DefaultA and SomMotB (Supplementary Tables 1 and 2). Thus, the observed network-level effects represent aggregate network tendencies rather than uniform responses across all constituent ROIs.

In summary, dynamic ALFF revealed non-monotonic, phase-dependent, and network-specific responses to acute severe hypoxia. These responses distinguished a reversible decompensation phase from earlier adaptation and were followed by a pronounced rebound after reoxygenation. Network-level differentiation, particularly between DefaultA and SomMotB, further indicated that severe hypoxia does not produce uniform suppression of spontaneous BOLD activity. Together, these findings suggest that dALFF captures a distinct aspect of the brain’s response to metabolic stress and may provide a useful measure for characterizing reversible functional state transitions and network-level reorganization during acute hypoxia.

## 6. Acknowledgements

The authors acknowledge the contributions of the Human Integrative Physiology Laboratory, the Clinical Research and Trials Unit, and the Special Purpose Processor Development Group at Mayo Clinic. The authors also thank Dr. Max Trenerry for early discussions related on the neuropsychological aspects of the study.

This work was supported by the Office of Naval Research and the National Institute of Biomedical Imaging and Bioengineering of the National Institutes of Health. The opinions, findings, and conclusions or recommendations expressed in this material are those of the authors and do not necessarily reflect the views of the Office of Naval Research.

## 7. Author contributions: CRediT

D.K. conceived the study, developed the analytical framework, performed the formal analysis, generated the visualizations, and wrote the original draft of the manuscript. K.U., C.R.H., N.G.C., D.R.H., M.J.J., T.B.C., and J.H. contributed to conceptualization. D.K., K.U., C.R.H., N.G.C., M.-H.I., J.D.T., K.M.W., C.C.W., J.W.S., S.E.B., M.A.B., D.R.H., M.J.J., T.B.C., and Y.S. contributed to methodology. E.M.G. contributed to investigation. C.R.H. and T.B.C. contributed to funding acquisition. T.B.C. and J.H. contributed to project administration. All authors contributed to review and editing of the manuscript and approved the final version.

## 8. Ethics approval statements

The study was conducted under an Institutional Review Board–approved protocol (Mayo Clinic IRB#16-004354). Experimental procedures were performed in accordance with the ethical standards set by the *Declaration of Helsinki*. All participants provided written informed consent prior to participation.

## 11. Declaration of generative AI and AI-assisted technologies in the manuscript preparation process

During the preparation of this work the authors used ChatGPT (OpenAI) in order to improve language and readability of the manuscript. After using this tool, the authors reviewed and edited the content as needed and take full responsibility for the content of the published article.

## 12. Data availability

The raw data generated in this study are not publicly available because of restrictions related to participant privacy and institutional policies.

## References

Achuthan, S.K., Coburn, K.L., Beckerson, M.E., Kana, R.K., 2023. Amplitude of low frequency fluctuations during resting state fMRI in autistic children. Autism Research 16, 84–98.

Behzadi, Y., Restom, K., Liau, J., Liu, T.T., 2007. A component based noise correction method (CompCor) for BOLD and perfusion based fMRI. NeuroImage 37, 90–101.

Bengtsson, J., Bake, B., Johansson, Å., Bengtson, J.P., 2001. End-tidal to arterial oxygen tension difference as an oxygenation index. Acta Anaesthesiologica Scandinavica 45, 357–363.

Birn, R.M., 2012. The role of physiological noise in resting-state functional connectivity. NeuroImage 62, 864–870.

Birn, R.M., Smith, M.A., Jones, T.B., Bandettini, P.A., 2008. The respiration response function: the temporal dynamics of fMRI signal fluctuations related to changes in respiration. NeuroImage 40, 644–654.

Biswal, B.B., Uddin, L.Q., 2025. The history and future of resting-state functional magnetic resonance imaging. Nature 641, 1121–1131.

Bloomfield, P.M., Connell, C.J.W., Shaw, D.M., Fan, J.L., Fisher, J.P., Gant, N., 2026. Systematic review and meta-analysis of acute hypoxia and cognition: Domain-specific effects and the role of barometric pressure and carbon dioxide. Neurosci Biobehav Rev 184, 106603.

Brugniaux, J.V., Hodges, A.N., Hanly, P.J., Poulin, M.J., 2007. Cerebrovascular responses to altitude. Respir Physiol Neurobiol 158, 212–223.

Chen, X.Q., Dong, J., Niu, C.Y., Fan, J.M., Du, J.Z., 2007. Effects of hypoxia on glucose, insulin, glucagon, and modulation by corticotropin-releasing factor receptor type 1 in the rat. Endocrinology 148, 3271–3278.

Cox, R.W., 1996. AFNI: software for analysis and visualization of functional magnetic resonance neuroimages. Comput Biomed Res 29, 162–173.

Craig, A.D., 2009. How do you feel--now? The anterior insula and human awareness. Nat Rev Neurosci 10, 59–70.

Delli Pizzi, S., Gambi, F., Di Pietro, M., Caulo, M., Sensi, S.L., Ferretti, A., 2023. BOLD cardiorespiratory pulsatility in the brain: from noise to signal of interest. Front Hum Neurosci 17, 1327276.

Di, X., Biswal, B.B., 2015. Dynamic brain functional connectivity modulated by resting-state networks. Brain Structure & Function 220, 37–46.

Foo, T.K.F., Laskaris, E., Vermilyea, M., Xu, M., Thompson, P., Conte, G., Van Epps, C., Immer, C., Lee, S.K., Tan, E.T., Graziani, D., Mathieu, J.B., Hardy, C.J., Schenck, J.F., Fiveland, E., Stautner, W., Ricci, J., Piel, J., Park, K., Hua, Y., Bai, Y., Kagan, A., Stanley, D., Weavers, P.T., Gray, E., Shu, Y., Frick, M.A., Campeau, N.G., Trzasko, J., Huston, J., 3rd, Bernstein, M.A., 2018. Lightweight, compact, and high-performance 3T MR system for imaging the brain and extremities. Magn Reson Med 80, 2232–2245.

Fu, Z., Tu, Y., Di, X., Du, Y., Pearlson, G.D., Turner, J.A., Biswal, B.B., Zhang, Z., Calhoun, V.D., 2018. Characterizing dynamic amplitude of low-frequency fluctuation and its relationship with dynamic functional connectivity: An application to schizophrenia. NeuroImage 180, 619–631.

Glover, G.H., Li, T.Q., Ress, D., 2000. Image-based method for retrospective correction of physiological motion effects in fMRI: RETROICOR. Magnetic Resonance in Medicine 44, 162–167.

Gong, H., Liu, Y.X., Xiaoluo, Q.L., Gong, M.F., Liu, Z., Wu, S.R., Chen, Z.Y., Liu, T.Y., Zhao, J.H., Wang, L., Fan, X.T., Xu, H.W., 2025. Unraveling the complexity of cognitive impairment following high-altitude exposure: from preclinical animal models to human organoids. Front Neurosci 19, 1679858.

Grzybowska-Ganszczyk, D., Nowak, Z., Opara, J.A., Nowak-Lis, A., 2025. Hypoxia and Cognitive Functions in Patients Suffering from Cardiac Diseases: A Narrative Review. J Clin Med 14.

Han, Y., Wang, J., Zhao, Z., Min, B., Lu, J., Li, K., He, Y., Jia, J., 2011. Frequency-dependent changes in the amplitude of low-frequency fluctuations in amnestic mild cognitive impairment: a resting-state fMRI study. NeuroImage 55, 287–295.

Harris, A.D., Murphy, K., Diaz, C.M., Saxena, N., Hall, J.E., Liu, T.T., Wise, R.G., 2013. Cerebral blood flow response to acute hypoxic hypoxia. NMR Biomed 26, 1844–1852.

Hofmeijer, J., Mulder, A.T., Farinha, A.C., van Putten, M.J., le Feber, J., 2014. Mild hypoxia affects synaptic connectivity in cultured neuronal networks. Brain Res 1557, 180–189.

Jo, H.J., Saad, Z.S., Simmons, W.K., Milbury, L.A., Cox, R.W., 2010. Mapping sources of correlation in resting state FMRI, with artifact detection and removal. NeuroImage 52, 571–582.

Kang, D., Uchida, K., Haider, C.R., Campeau, N.G., In, M.H., Gray, E.M., Trzasko, J.D., Welker, K.M., Bernstein, M.A., Trenerry, M.R., Holmes Iii, D.R., Joyner, M.J., Curry, T.B., Huston Iii, J., Shu, Y., 2026. Brain functional connectivity initiates structured reorganization at a critical oxygen threshold during hypoxia. Brain Res Bull 235, 111748.

Kang, D., Uchida, K., Haider, C.R., Campeau, N.G., In, M.H., Gray, E.M., Trzasko, J.D., Welker, K.M., Gunter, J.L., Bernstein, M.A., Trenerry, M.R., Holmes, D.R., 3rd, Joyner, M.J., Curry, T.B., Huston, J., 3rd, Shu, Y., 2025. Regional variation in cerebral oxygen metabolism during acute severe hypoxia with temporary cognitive impairment. NeuroImage 316, 121302.

Mascarenhas, A., Braga, A., Majernikova, S.M., Nizari, S., Marletta, D., Theparambil, S.M., Aziz, Q., Marina, N., Gourine, A.V., 2025. On the mechanisms of brain blood flow regulation during hypoxia. J Physiol 603, 2263–2280.

McMorris, T., Hale, B.J., Barwood, M., Costello, J., Corbett, J., 2017. Effect of acute hypoxia on cognition: A systematic review and meta-regression analysis. Neurosci Biobehav Rev 74, 225–232.

Naismith, S.L., Duffy, S.L., Cross, N., Grunstein, R., Terpening, Z., Hoyos, C., D’Rozario, A., Lagopoulos, J., Osorio, R.S., Shine, J.M., McKinnon, A.C., 2020. Nocturnal Hypoxemia Is Associated with Altered Parahippocampal Functional Brain Connectivity in Older Adults at Risk for Dementia. J Alzheimers Dis 73, 571–584.

Ogoh, S., 2015. Cerebral blood flow regulation during hypoxia. Exp Physiol 100, 109–110.

Ortiz-Prado, E., Dunn, J.F., Vasconez, J., Castillo, D., Viscor, G., 2019. Partial pressure of oxygen in the human body: a general review. Am J Blood Res 9, 1–14.

Phillips, J.B., Horning, D., Funke, M.E., 2015. Cognitive and perceptual deficits of normobaric hypoxia and the time course to performance recovery. Aerosp Med Hum Perform 86, 357–365.

Post, T.E., Heijn, L.G., Jordan, J., van Gerven, J.M.A., 2023. Sensitivity of cognitive function tests to acute hypoxia in healthy subjects: a systematic literature review. Front Physiol 14, 1244279.

Raichle, M.E., 2015. The brain’s default mode network. Annu Rev Neurosci 38, 433–447.

Raichle, M.E., Gusnard, D.A., 2002. Appraising the brain’s energy budget. Proc Natl Acad Sci U S A 99, 10237–10239.

Reuter, M., Schmansky, N.J., Rosas, H.D., Fischl, B., 2012. Within-subject template estimation for unbiased longitudinal image analysis. NeuroImage 61, 1402–1418.

Schaefer, A., Kong, R., Gordon, E.M., Laumann, T.O., Zuo, X.N., Holmes, A.J., Eickhoff, S.B., Yeo, B.T.T., 2018. Local-Global Parcellation of the Human Cerebral Cortex from Intrinsic Functional Connectivity MRI. Cereb Cortex 28, 3095–3114.

Tao, S., Trzasko, J.D., Gunter, J.L., Weavers, P.T., Shu, Y., Huston, J., Lee, S.K., Tan, E.T., Bernstein, M.A., 2017. Gradient nonlinearity calibration and correction for a compact, asymmetric magnetic resonance imaging gradient system. Phys Med Biol 62, N18–N31.

Taylor, L., Watkins, S.L., Marshall, H., Dascombe, B.J., Foster, J., 2015. The Impact of Different Environmental Conditions on Cognitive Function: A Focused Review. Front Physiol 6, 372.

Todd, M.M., Wu, B., Maktabi, M., Hindman, B.J., Warner, D.S., 1994. Cerebral blood flow and oxygen delivery during hypoxemia and hemodilution: role of arterial oxygen content. Am J Physiol 267, H2025–2031.

Tommasin, S., Mascali, D., Gili, T., Assan, I.E., Moraschi, M., Fratini, M., Wises, R.G., Macaluso, E., Mangia, S., Giove, F., 2017. Task-Related Modulations of BOLD Low-Frequency Fluctuations within the Default Mode Network. Frontiers in Physics 5.

Turner, C.E., Barker-Collo, S.L., Connell, C.J., Gant, N., 2015. Acute hypoxic gas breathing severely impairs cognition and task learning in humans. Physiol Behav 142, 104–110.

Turner, R., Rasmussen, P., Gatterer, H., Tremblay, J.C., Roche, J., Strapazzon, G., Roveri, G., Lawley, J., Siebenmann, C., 2024. Cerebral blood flow regulation in hypobaric hypoxia: role of haemoconcentration. J Physiol 602, 5643–5657.

Uchida, K., Baker, S.E., Wiggins, C.C., Senefeld, J.W., Shepherd, J.R.A., Trenerry, M.R., Buchholtz, Z.A., Clifton, H.R., Holmes, D.R., Joyner, M.J., Curry, T.B., 2020a. A Novel Method to Measure Transient Impairments in Cognitive Function During Acute Bouts of Hypoxia. Aerosp Med Hum Perform 91, 839–844.

Uchida, K., Baker, S.E., Wiggins, C.C., Senefeld, J.W., Shepherd, J.R.A., Trenerry, M.R., Clifton, H.R., Holmes, D.R., Joyner, M.J., Curry, T.B., 2020b. Relationship between Decreased Oxygenation during Acute Hypoxia and Cognitive Deterioration in Healthy Humans. The FASEB Journal 34, 1–1.

Wang, L., Liu, Q., Zheng, Z., Chen, W., Shen, Y., Li, T., Feng, C., 2025. Common neural dysfunction in psychiatric disorders: Insights from a meta-analysis of resting-state fMRI studies. Transl Psychiatry 16, 29.

Wang, X., Cui, L., Ji, X., 2022. Cognitive impairment caused by hypoxia: from clinical evidences to molecular mechanisms. Metab Brain Dis 37, 51–66.

Yang, H., Long, X.Y., Yang, Y.H., Yan, H., Zhu, C.Z., Zhou, X.P., Zang, Y.F., Gong, Q.Y., 2007. Amplitude of low frequency fluctuation within visual areas revealed by resting-state functional MRI. NeuroImage 36, 144–152.

Yeo, B.T., Krienen, F.M., Sepulcre, J., Sabuncu, M.R., Lashkari, D., Hollinshead, M., Roffman, J.L., Smoller, J.W., Zollei, L., Polimeni, J.R., Fischl, B., Liu, H., Buckner, R.L., 2011. The organization of the human cerebral cortex estimated by intrinsic functional connectivity. J Neurophysiol 106, 1125–1165.

Zheng, R., Chen, Y., Jiang, Y., Wen, M., Zhou, B., Li, S., Wei, Y., Yang, Z., Wang, C., Cheng, J., Zhang, Y., Han, S., 2021. Dynamic Altered Amplitude of Low-Frequency Fluctuations in Patients With Major Depressive Disorder. Front Psychiatry 12, 683610.

