## Supplementary Tables and Figures for "Dynamic ALFF Reveals Phase- and Network-Dependent Responses During Acute Severe Hypoxia"

### 1. Supplementary Tables

**Supplementary Table 1.** ROI-level dALFF changes within DefaultA during the decompensation phase.

| Networks | Hemisphere | ROI label | Median ΔALFF at t = 360 s (%) | IQR (%) |
|---|---|---|---|---|
| DefaultA | LH | pCunPCC_1 | -44.6 | [-52.6, -28.0] |
| DefaultA | LH | IPL_2 | -41.3 | [-56.1, -26.6] |
| DefaultA | LH | pCunPCC_2 | -37.0 | [-49.3, -20.9] |
| DefaultA | RH | pCunPCC_1 | -35.4 | [-52.5, -24.4] |
| DefaultA | LH | pCunPCC_3 | -34.8 | [-51.4, -4.3] |
| DefaultA | RH | pCunPCC_2 | -33.5 | [-40.9, -23.6] |
| DefaultA | LH | IPL_1 | -33.1 | [-43.1, -8.3] |
| DefaultA | RH | pCunPCC_5 | -32.3 | [-41.3, -15.7] |
| DefaultA | RH | IPL_2 | -31.6 | [-45.7, -9.1] |
| DefaultA | RH | PFCd_2 | -27.4 | [-37.4, -19.1] |
| DefaultA | LH | pCunPCC_6 | -26.7 | [-45.2, -8.6] |
| DefaultA | LH | PFCd_3 | -26.1 | [-35.7, -16.1] |
| DefaultA | LH | PFCd_1 | -24.7 | [-35.8, -15.4] |
| DefaultA | RH | IPL_1 | -24.6 | [-45.6, -13.9] |
| DefaultA | RH | PFCm_4 | -23.5 | [-32.9, 10.7] |
| DefaultA | LH | pCunPCC_7 | -23.4 | [-39.8, -2.8] |
| DefaultA | LH | PFCm_3 | -23.2 | [-41.8, 11.3] |
| DefaultA | LH | pCunPCC_4 | -23.0 | [-28.9, 13.6] |
| DefaultA | LH | PFCd_2 | -22.9 | [-31.2, -11.2] |
| DefaultA | RH | PFCd_1 | -21.8 | [-44.0, 12.2] |
| DefaultA | LH | PFCm_2 | -20.4 | [-23.8, 21.7] |
| DefaultA | LH | PFCm_1 | -16.9 | [-39.4, 1.7] |
| DefaultA | LH | PFCm_4 | -15.6 | [-22.1, 1.4] |
| DefaultA | LH | PFCm_5 | -13.2 | [-43.7, 24.7] |
| DefaultA | RH | Temp_1 | -12.4 | [-27.3, 13.7] |
| DefaultA | RH | PFCm_1 | -10.0 | [-36.3, 5.9] |
| DefaultA | RH | PFCm_3 | -5.7 | [-24.4, 0.8] |
| DefaultA | RH | pCunPCC_3 | -4.3 | [-35.8, 6.3] |
| DefaultA | RH | PFCm_2 | -3.1 | [-36.1, 14.7] |

| | | | | |
|---|---|---|---|---|
| DefaultA | RH | pCunPCC_4 | -2.5 | [-22.8, 65.3] |
| DefaultA | LH | pCunPCC_5 | 3.5 | [-32.9, 69.5] |
| DefaultA | LH | PFCm_6 | 5.0 | [-3.4, 25.4] |
| DefaultA | RH | PFCm_5 | 5.8 | [-41.8, 19.7] |
| DefaultA | RH | PFCm_6 | 19.8 | [11.4, 34.3] |

**Note**. ROIs are ordered by median ΔALFF at t = 360 s in ascending order.
**Abbreviations**: LH, left hemisphere; RH, right hemisphere; pCunPCC, precuneus/posterior cingulate cortex; IPL, inferior parietal lobule; PFCd, dorsal prefrontal cortex; PFCm, medial prefrontal cortex; Temp, temporal cortex.

**Supplementary Table 2**. ROI-level dALFF changes within SomMotB during the decompensation phase.

| Networks | Hemisphere | ROI label | Median ΔALFF at t = 360 s (%) | IQR (%) |
|---|---|---|---|---|
| SomMotB | RH | Aud_2 | 36.0 | [-0.9, 51.7] |
| SomMotB | RH | Ins_1 | 34.5 | [-6.5, 50.0] |
| SomMotB | RH | S2_5 | 27.5 | [-16.0, 62.0] |
| SomMotB | RH | S2_4 | 24.3 | [-10.9, 54.8] |
| SomMotB | LH | S2_2 | 24.1 | [-14.9, 41.3] |
| SomMotB | RH | S2_2 | 23.4 | [-6.1, 43.5] |
| SomMotB | RH | S2_8 | 20.8 | [-11.7, 25.3] |
| SomMotB | LH | S2_3 | 19.6 | [-27.3, 43.6] |
| SomMotB | LH | S2_1 | 18.5 | [-27.8, 47.1] |
| SomMotB | LH | Aud_1 | 17.9 | [-8.4, 44.3] |
| SomMotB | LH | Aud_4 | 15.6 | [-3.8, 31.0] |
| SomMotB | RH | S2_1 | 15.0 | [-15.8, 55.0] |
| SomMotB | RH | Aud_1 | 12.9 | [-14.8, 52.0] |
| SomMotB | RH | S2_7 | 11.6 | [-12.0, 56.7] |
| SomMotB | LH | Ins_1 | 9.3 | [-23.1, 54.3] |
| SomMotB | LH | Aud_2 | 9.3 | [-15.7, 30.5] |
| SomMotB | RH | Aud_3 | 3.3 | [-5.6, 45.2] |
| SomMotB | RH | S2_3 | -0.1 | [-14.1, 75.3] |
| SomMotB | RH | Cent_2 | -2.5 | [-29.7, 36.4] |
| SomMotB | LH | Cent_3 | -5.6 | [-18.4, 16.5] |
| SomMotB | RH | S2_6 | -6.9 | [-24.0, 22.3] |
| SomMotB | LH | S2_4 | -6.9 | [-28.3, 13.0] |

| | | | | |
|---|---|---|---|---|
| SomMotB | LH | Cent_1 | -8.3 | [-21.5, 27.7] |
| SomMotB | LH | Cent_5 | -9.1 | [-20.5, 46.4] |
| SomMotB | LH | Aud_3 | -9.6 | [-29.0, 18.3] |
| SomMotB | LH | Cent_2 | -10.1 | [-28.8, 44.3] |
| SomMotB | LH | S2_5 | -11.0 | [-42.2, 23.6] |
| SomMotB | RH | Cent_1 | -14.3 | [-32.4, 31.9] |
| SomMotB | RH | Cent_3 | -14.6 | [-21.0, 11.9] |
| SomMotB | LH | S2_6 | -14.7 | [-44.4, 24.3] |
| SomMotB | LH | Cent_4 | -20.8 | [-39.9, 49.4] |

**Note**. ROIs are ordered by median ΔALFF at t = 360 s in descending order.
**Abbreviations**: LH, left hemisphere; RH, right hemisphere; Aud, auditory cortex; Ins, insular cortex; S2, secondary somatosensory cortex; Cent, central sensorimotor cortex.

#### 2. Supplementary Figures

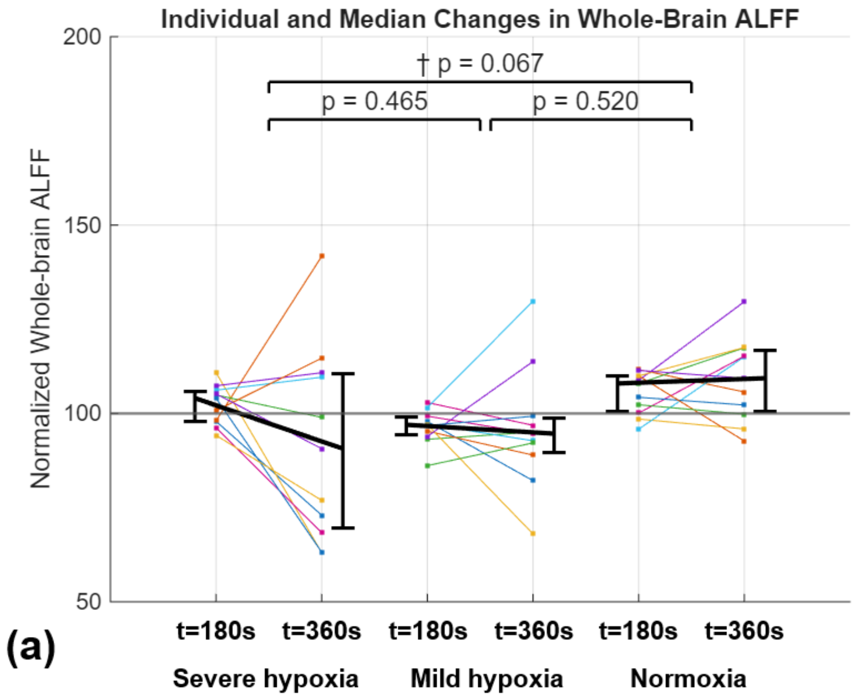


**Supplementary Figure 1. Individual and group-level changes in whole-brain ALFF relative to baseline across oxygen conditions.** Whole-brain amplitude of low-frequency fluctuations (ALFF), normalized to baseline, is shown at 180 s and 360 s during severe hypoxia, sustained mild hypoxia, and normoxia. Colored lines represent individual participants, while black lines and error bars indicate the group median and interquartile range (IQR), respectively. Whole-brain ALFF showed a greater decrease from baseline during severe hypoxia, whereas smaller changes were observed during mild hypoxia and normoxia. Pairwise comparisons of temporal changes showed no significant differences between severe and mild hypoxia ($p = 0.465$) or between mild hypoxia and normoxia ($p = 0.520$), while the severe hypoxia–normoxia comparison showed a trend toward significance (†$p = 0.067$). The horizontal gray line at 100 indicates the baseline-normalized reference level.

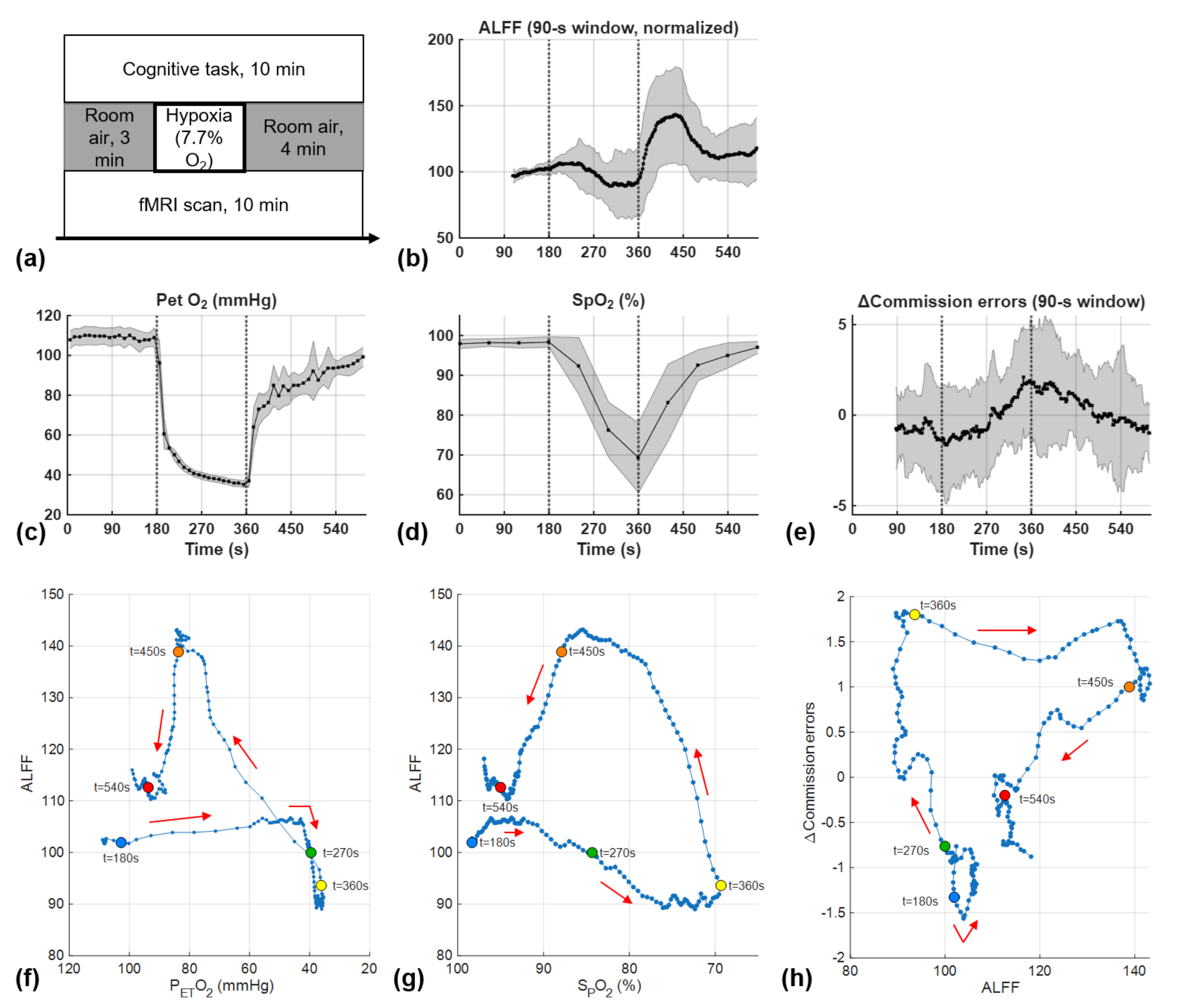


**Supplementary Figure 2. Dynamic ALFF trajectories during severe hypoxia.** (a) Experimental paradigm for the severe hypoxia condition. (b) Time-resolved whole-brain dynamic amplitude of low-frequency fluctuations (dALFF), calculated using a 90-s window and normalized to baseline. (c) End-tidal oxygen partial pressure (PETO2) and (d) peripheral oxygen saturation (SpO2) over time. (e) Time-resolved changes in Δcommission errors calculated using a 90-s window. (f) dALFF trajectory plotted against corresponding PETO2 values, (g) dALFF trajectory plotted against corresponding SpO2 values, and (h) Δcommission-error trajectory plotted against dALFF. Colored markers indicate selected time points (180, 270, 360, 450, and 540 s), and arrows indicate the temporal direction of the trajectories. Black lines indicate the group mean, and gray shaded regions indicate the standard deviation.

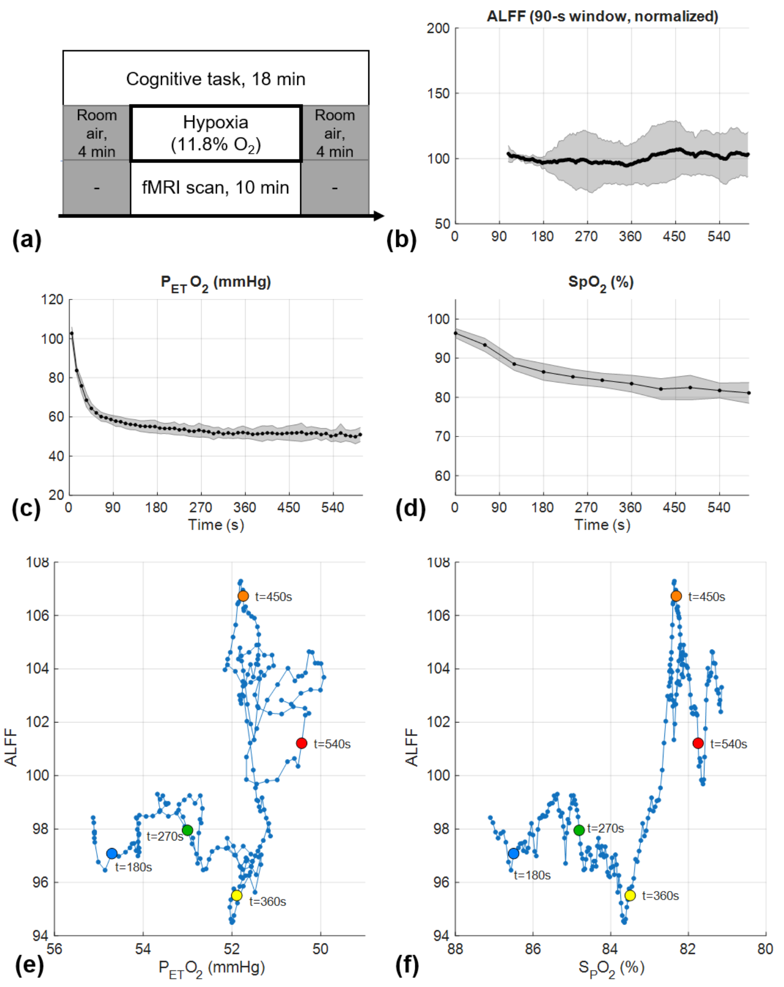


**Supplementary Figure 3. Dynamic ALFF trajectories during sustained mild hypoxia.** (a) Experimental paradigm for the sustained mild hypoxia condition. (b) Time-resolved whole-brain dynamic amplitude of low-frequency fluctuations (dALFF), calculated using a 90-s window and normalized to baseline. (c) End-tidal oxygen partial pressure (PETO2) and (d) peripheral oxygen saturation (SpO2) over time. (e) dALFF trajectory plotted against corresponding PETO2 values and (f) dALFF trajectory plotted against corresponding SpO2 values. Colored markers indicate selected time points (180, 270, 360, 450, and 540 s), illustrating the temporal evolution of dALFF relative to systemic oxygenation. Black lines indicate the group mean, and gray shaded regions indicate the standard deviation.

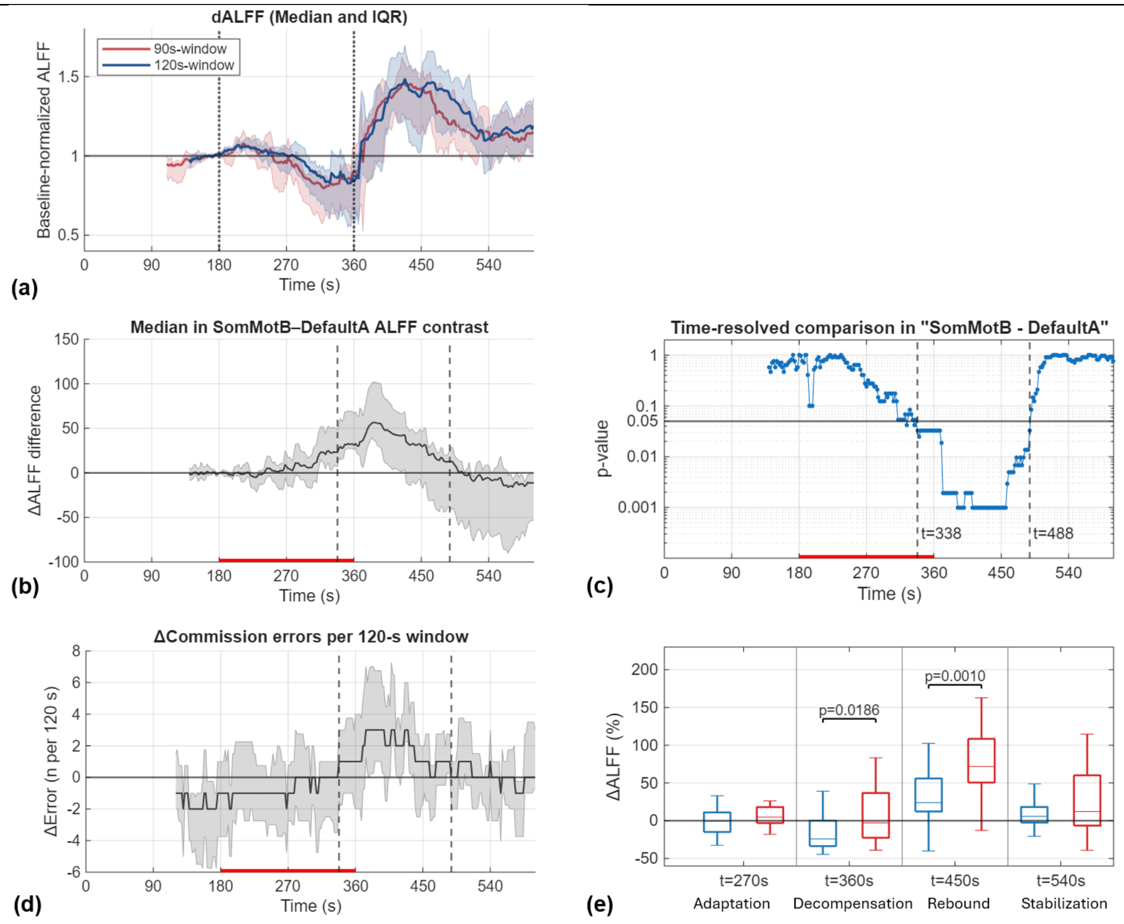


**Supplementary Figure 4. Robustness of dALFF dynamics and network-level differentiation using a 120-s sliding window.** (a) Comparison of whole-brain dALFF trajectories obtained using 90-s and 120-s causal sliding windows. Solid lines indicate medians and shaded regions indicate interquartile ranges (IQRs). (b) Median SomMotB–DefaultA dALFF contrast across subjects using the 120-s window. (c) Time-resolved exact paired Wilcoxon signed-rank comparisons between SomMotB and DefaultA. The horizontal line indicates the nominal threshold of $p = 0.05$. Continuous nominal differentiation was observed from approximately 338 to 488 s. Because adjacent windows overlapped and the time-resolved tests were not corrected for multiple comparisons, this interval should be considered exploratory. (d) Median Δcommission errors across subjects calculated using 120-s sliding windows. The period of increased commission errors broadly overlapped with the interval of network differentiation. (e) Distributions of ΔALFF in DefaultA and SomMotB at the same representative time points used in the primary 90-s analysis: adaptation (t = 270 s), decompensation (t = 360 s), rebound (t = 450 s), and stabilization (t = 540 s). Exact paired Wilcoxon signed-rank tests showed significant differences during decompensation and rebound. Displayed $p$ values are unadjusted. Red bars denote the hypoxia exposure period.

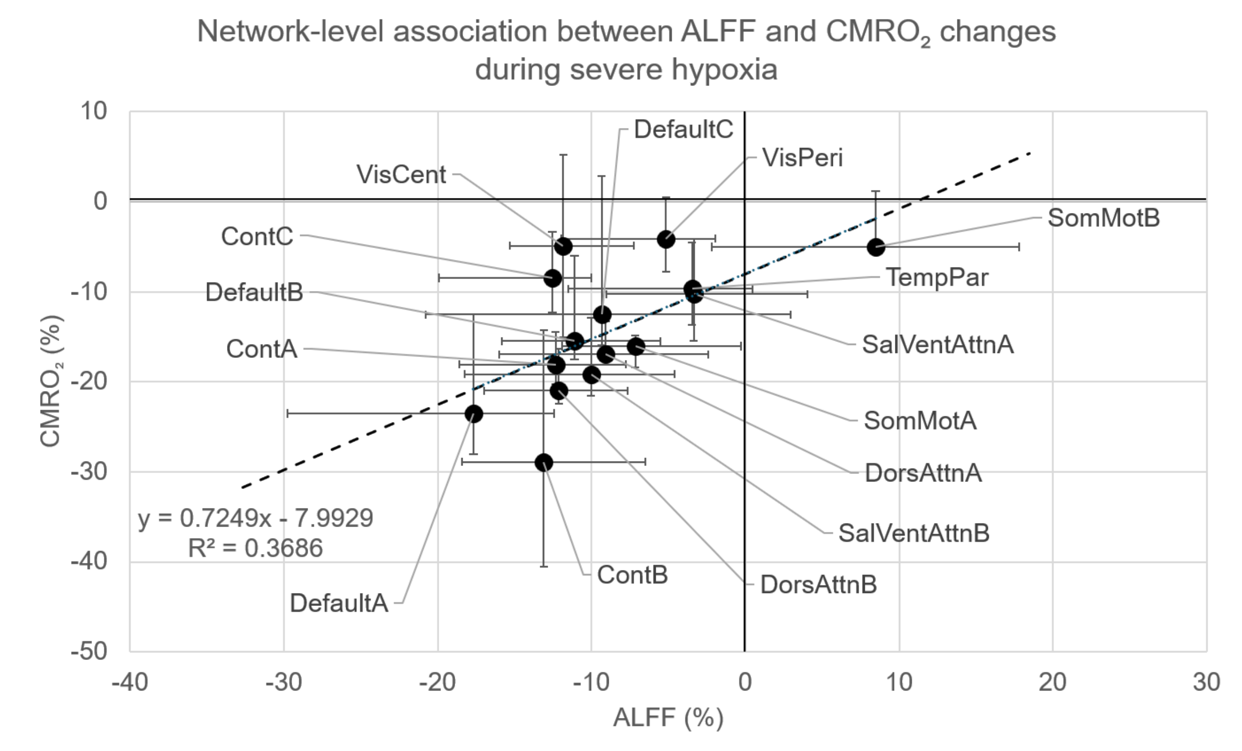


**Supplementary Figure 5. Network-level associations between ALFF and CMRO2 changes during severe hypoxia.** Scatter plot showing network-wise changes in ALFF and estimated CMRO2 at the decompensation phase (t ~ 350 s). Each point represents the mean change within a functional network relative to the baseline period (<180 s). Error bars indicate the interquartile range (Q1–Q3) across ROIs within each network. Estimated CMRO2 values were derived using the modified Davis model described in our previous study (Kang et al., 2025) and aggregated at the network level for comparison with ALFF changes. Across networks, reductions in ALFF were moderately associated with decreases in estimated CMRO2 (linear fit shown; R2 = 0.37). Networks showing larger suppression of local BOLD signal fluctuations tended to exhibit greater metabolic reductions. This exploratory analysis suggests that network-level decreases in ALFF may reflect reduced metabolic demand during severe hypoxia.
